# Analysis of high-molecular-weight proteins using MALDI-TOF MS and Machine Learning for the differentiation of clinically relevant *Clostridioides difficile* ribotypes

**DOI:** 10.1101/2024.06.18.599569

**Authors:** Ana Candela, David Rodriguez-Temporal, Mario Blázquez-Sánchez, Manuel J. Arroyo, Mercedes Marín, Luis Alcalá, Germán Bou, Belén Rodríguez-Sánchez, Marina Oviaño

**Affiliations:** Clinical Microbiology Department, Complexo Hospitalario Universitario A Coruña, Institute of Biomedical Research A Coruña (INIBIC) A Coruña, Spain; Clinical Microbiology and Infectious Diseases Department, Hospital General Universitario Gregorio Marañón and Institute of Health Research Gregorio Marañón (IiSGM) Madrid, Spain; Clover Bioanalytical Software, Av. del Conocimiento, 41, 18016 Granada, Spain; CIBER de Enfermedades Respiratorias (CIBERES CB06/06/0058), Madrid 28007, Spain; CIBER de Enfermedades Infecciosas (CIBERINFEC CB21/13/00055)

**Keywords:** *Clostridioides difficile*, MALDI-TOF MS, Machine Learning, Ribotyping, MALDI-TOF, high-molecular-weight proteins, *Clostridioides difficile*, Machine Learning, ribotyping

## Abstract

*Clostridioides difficile* is the main cause of antibiotic related diarrhea and some ribotypes (RT), such as RT027, RT181 or RT078, are considered high risk clones. A fast and reliable approach for *C. difficile* ribotyping is needed for a correct clinical approach. This study analyses high-molecular-weight proteins for *C. difficile* ribotyping with MALDI-TOF MS. Sixty-nine isolates representative of the most common ribotypes in Europe were analyzed in the 17,000-65,000 *m/z* region and classified into 4 categories (RT027, RT181, RT078 and ‘Other RTs’). Five supervised Machine Learning algorithms were tested for this purpose: K-Nearest Neighbors, Support Vector Machine, Partial Least Squares-Discriminant Analysis, Random Forest and Light-Gradient Boosting Machine. All algorithms yielded cross-validation results >70%, being RF and Light-GBM the best performing, with 88% of agreement. Area under the ROC curve of these two algorithms was >0.9. RT078 was correctly classified with 100% accuracy and isolates from the RT181 category could not be differentiated from RT027.

## Introduction

*Clostridioides difficile* is an anaerobic Gram-positive rod that can persist on surfaces and in the environment, and is resistant to most conventional disinfectants such as alcohol or chlorhexidine due to its ability to form spores. This makes it highly transmissible if good hygiene and infection control measures are not implemented (1). This microorganism is the main cause of antibiotic related diarrhea and represents a public health concern due to its high morbidity and mortality rates and its involvement in nosocomial outbreaks.

Some *C. difficile* ribotypes (RTs) have shown to be more virulent and/or involved in nosocomial outbreaks due to the production of toxins. *C. difficile* RT NAP1/B1/RT027 and the recently described RT181 (“027-like”) (2-4) also present a deletion of 1 bp at position 117 of the regulatory gene *tcdC*, a pathogenicity locus that downregulates the production of toxins. The consequence of this deletion is the hyperproduction of toxins, which makes these RTs more pathogenic and associated to more severe outcomes (5-7).

*C. difficile* characterization and ribotyping should be fast and reliable, to enable the correct antibiotic implementation and infection control measures. Gold Standard techniques in the USA and Europe are Pulsed Field Gel Electrophoresis (PFGE) and PCR Ribotyping, respectively (8). These techniques are laborious and require specialized personnel trained in molecular biology. Also, they are cumbersome and final results are obtained after several days. Among the commercially available molecular techniques, one of the most used is Xpert® *C. difficile* BT (Cepheid, Sunnyvale, CA, USA), which allows the detection of toxin B, the binary toxin and the deletion at position 117 in gene *tcdC* (9). However, it has been shown that this test does not distinguish the different toxigenic RTs that host the deletion in gene *tcdC* (2, 10, 11).

Matrix-Assisted Laser Desorption/Ionization Time of Flight Mass Spectrometry (MALDI-TOF MS) represents an alternative to the previously described reference methods since it is a technology available in most microbiology laboratories nowadays, easy to use and with a time-around time of only some minutes. Apart from the initial investment for the acquisition of the instrument, the cost per sample is lower than that of molecular techniques. Besides identification, MALDI-TOF MS has been applied for bacterial typing and antibiotic resistance detection in several species like *Pseudomonas aeruginosa, Staphylococcus aureus, Streptococcus pneumoniae* or *Escherichia coli* (12-15). The default region for spectra analysis is between 2,000 and 20,000 *m/z* -where most ribosomal proteins are found-, although it can be modified to analyze a higher molecular weight range.

For *C. difficile*, few studies have evaluated MALDI-TOF MS as a ribotyping alternative, and even fewer analyzed high molecular weight proteins. *C. difficile* external surface layer (S-layer) is composed of proteins with a molecular weight ranging from 40 to 200 kDa (16, 17). The study of this mass region increases the number of peaks available, expanding the chances for differentiation among different isolates. The aim of this study was to develop a rapid MALDI-TOF MS method for the differentiation of hypervirulent *C. difficile* RTs based the analysis of high molecular weight proteins.

## Materials and Methods

### Bacterial isolates and molecular characterization

A total of n=69 *C. difficile* isolates representing the most prevalent RTs in Spain and Europe were included in this study. All strains were isolated from clinical stool samples in Spain and ribotyped at Hospital General Universitario Gregorio Marañón, in Madrid, Spain. Clinical samples were directly analyzed upon reception with the commercial PCR Xpert^®^ *C. difficile* BT (Cepheid, Sunnyvale, CA, USA) and cultured in CLO agar (Beckton Dickinson®, Franklin Lakes, NJ, US) for *C. difficile* isolation and ribotyping. After bacterial growth, isolates were identified by MALDI-TOF MS in an MBT Smart MALDI Biotyper (Bruker Daltonics, Bremen, Germany) with the updated database containing 11,096 mass spectra profiles. The standard on-plate protein extraction was applied with 1µl 100% formic acid followed by 1µl HCCA (α-Cyano-4-hydroxycinnamic acid) matrix solution.

Ribotyping was performed by PCR amplification of the intergenic spacer region between the 16S rRNA and the 23S rRNA followed by capillary electrophoresis. The results of this sequencing were interpreted with Bionumerics 5.0 software (bioMérieux, Marcy l’Etoile, France) (18-21). Isolates included in this study belonged to the following RTs (Table S1): RT027 (n=29), RT181 (n=7), RT078 (n=21) and n=12 strains from other less toxigenic RTs (RT001, RT106, RT207, RT014 and RT023).

### MALDI-TOF MS spectra acquisition

For spectra acquisition, isolates were plated on Schaedler agar (Beckton Dickinson®, Franklin Lakes, NJ, US) and incubated at 37°C in anaerobic atmosphere for 48h. A few colonies were spotted on the MALDI target plate in duplicates and overlaid with 1µl of trans-ferulic acid matrix at a concentration of 15 mg/ml and dissolved in a solution of 33% acetonitrile, 17% formic acid and 50% water, as described previously (22, 23). Trans-ferulic acid matrix forms crystals upon drying (Figure 1) and allows the analysis of proteins in a higher molecular weight region, yielding larger mass peaks and higher signal-to-baseline ratios in comparison with other organic matrices (22).

**Figure 1.**
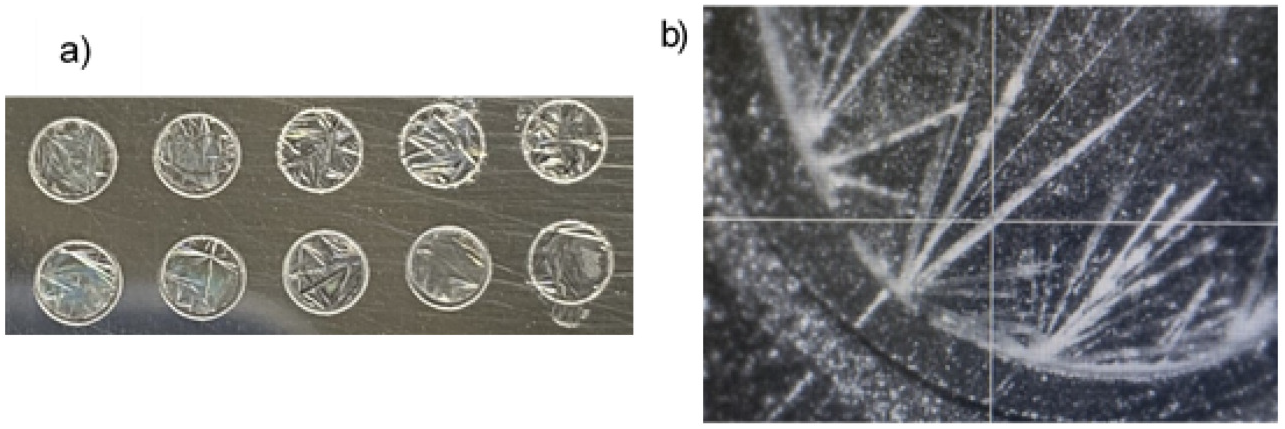
a) Example of crystallization of Trans-ferulic acid matrix on the MALDI plate; b) Zoomed-in portion of a MALDI spot during spectra acquisition.

Spectra were obtained manually in linear positive mode, reading over the formed crystals and acquired in the region 17,000-65,000 *m/z*, in pulses of 200 shots, for a summation of 800 spectra per strain and spot (Detector Gain 2.700V-2.800V, laser intensity 90%). Spectra were calibrated with the commercial calibrator PSII (Protein Standard II, Bruker Daltonics, Bremen, Germany).

### MALDI-TOF MS spectra analysis and data modeling

High molecular weight spectra were analyzed with the commercial platform Clover MS Data Analysis Software (MSDAS, https://www.clovermsdataanalysis.com; Clover Bioanalytical Software, Granada, Spain). Spectra were processed by a pipeline of: a) baseline subtraction using Top-Hat filter (factor 0.02), b) smoothing via Savitzky-Golay filter (window length: 11; polynomial order: 3), c) peak alignment with constant mass tolerance of 3 Da and linear mass tolerance of 600 ppm and d) Total Ion Current (TIC) normalization. Peaks were aligned in the 17,000-65,000 *m/z* region of the spectra and then merged in an average spectrum for each isolate. Full spectra were studied for this purpose.

Initial approach was to perform unsupervised algorithms to study the feasibility of using MALDI-TOF for the differentiation of *C. difficile* RTs analyzing the high molecular weight region. For that aim, hierarchical clustering with Principal Component Analysis (PCA) was performed. All isolates were included in this initial model and their high molecular weight spectra were compared.

After this initial study, several well-known supervised classification algorithms were applied: Random Forest (RF), Light-Gradient Boosting Machine (Light-GBM), Partial Least Squares-Discriminant Analysis (PLS-DA), Support Vector Machine (SVM) and K-Nearest Neighbor (KNN). These algorithms were trained by using a k-fold cross validation (k=5) as internal validation, meaning that 20% of the samples were randomly extracted from the model and blindly validated against it, on 5 different times. Hyperparameters applied for the construction of the models are summarized on Table S2.

Area under the receiver operating characteristic (AUROC) curve was obtained from these algorithms to evaluate their discrimination power for each category. The AUROC measures the ability of the model to discriminate among classes (i.e., higher values indicate greater ability) (24). A peak matrix with the full spectrum of each isolate (from 17,000 to 65,000 *m/z*) was employed as training data set. Biomarker analysis was applied to identify putative biomarkers capable of discriminating each category. For this purpose, ANOVA analysis and AUROC were calculated for all peaks found using a threshold of 1% of maximum intensity for peak selection.

## Results

All isolates were identified as *C. difficile* by MALDI-TOF MS with a score >2.0. Initial study by unsupervised methods showed three main groups in hierarchical clustering. First, a clear clustering of RT078, a second one composed of mainly isolates from the “Other RTs” category, and a third cluster that grouped together most isolates from categories RT027 and RT181 (Figure 2).

**Figure 2.**
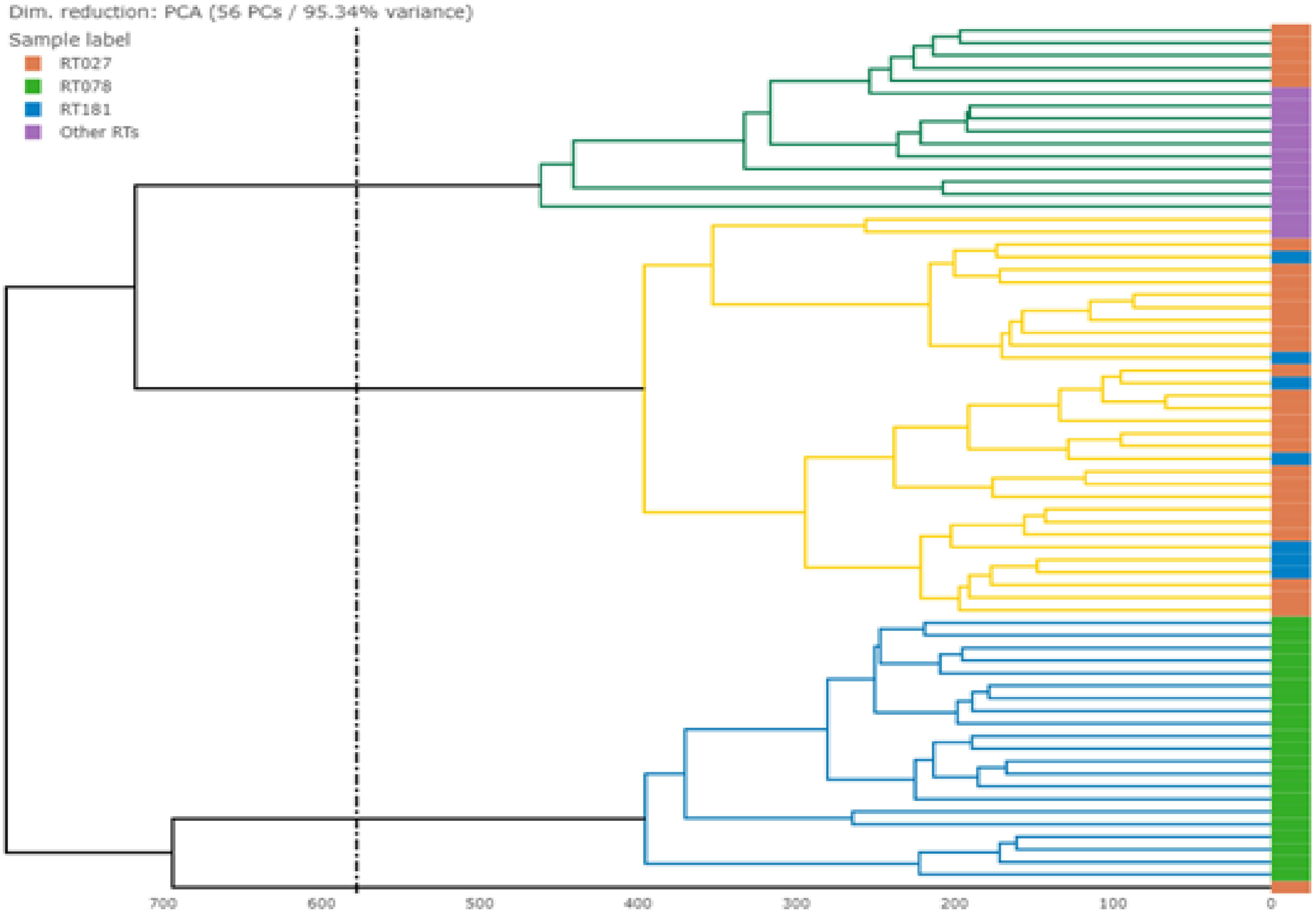
Dendrogram built with all isolates using Euclidean distance and Ward metric. Three main clusters could be observed. Dimensionality reduction was automatically applied to reach 95% of variance. In this experiment, there were automatically included 56 principal components.

All supervised algorithms evaluated yielded accuracy greater than 75%. The highest accuracy results for RT differentiation were obtained with the algorithms RF and Light-GBM. 5-fold cross validation of these two models yielded an accuracy of almost 90%. Differentiation of RT027 was possible with all algorithms with an accuracy >70% and a Positive Predictive Value (PPV) of 80.0% and 86.7% for RF and Light-GBM, respectively. RT078 could clearly be separated from the rest of RTs with an accuracy of 100% with all algorithms, and a PPV of 100% for RF and 95.5% for Light-GBM, with one RT027 isolate classified as RT078. Separation of RT181 from RT027 could not be achieved. Finally, “Other RTs” category also obtained an accuracy greater than 90% with RF and Light-GBM, and a PPV of 92.3% and 100%, respectively. Results of the 5-fold cross validation and distance plots of the algorithms are summarized in Table 1 and Figure 3, respectively.

**Table 1.** 5-fold cross validation of the different algorithms studied, showing the accuracy for each one of the categories of the model and the total accuracy.

| **% Correct** | **Number of isolates** | **KNN** | **PCA-SVM** | **Light-GBM** | **PLS-DA** | **RF** |
| --- | --- | --- | --- | --- | --- | --- |
| **RT027** | n=29 | 79.3% | 72.4% | 89.7% | 96.5% | 96.5% |
| **RT078** | n=21 | 100% | 100% | 100% | 100% | 100% |
| **RT181** | n=7 | 0% | 57.1% | 42.9% | 0% | 0% |
| **Other RTs** | n=12 | 66.7% | 83.3% | 91.7% | 83.3% | 100% |
| **Total % Correct** | n=69 | 75.4% | 81.2% | **88.4%** | 85.5% | **88.4%** |

KNN: K-Nearest Neighbor; PCA-SVM: Principal Component Analysis-Support Vector Machine; Light-GBM: Light Gradient Boosting Machine; PLS-DA: Partial Least Squares Discriminant Analysis; RF: Random Forest.

**Figure 3.**
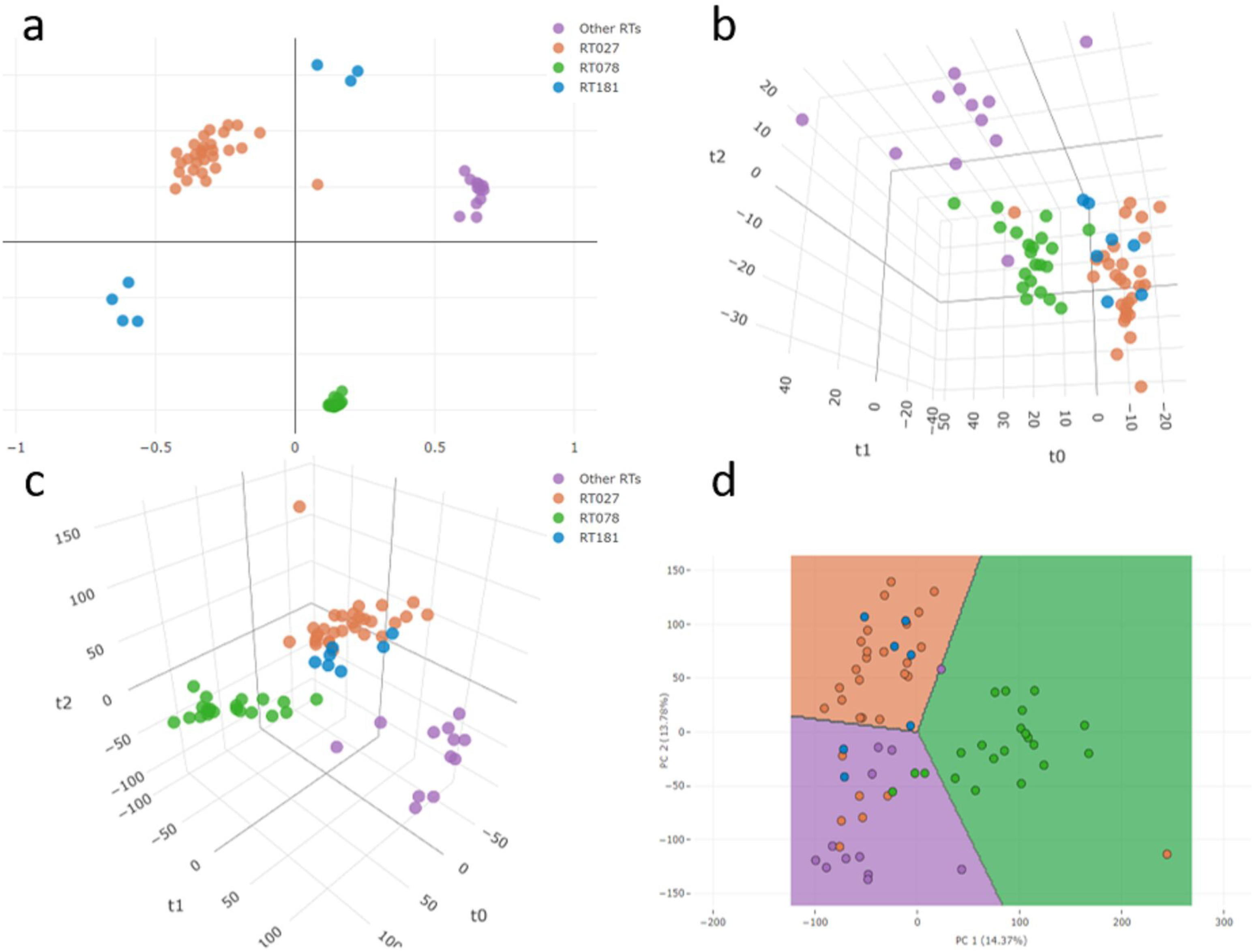
Distance plots of the algorithms studied: a) Random Forest; b) K-Nearest Neighbor; c) Partial Least Squares-Discriminant Analysis; d) Support Vector Machine.

AUROC was greater than 0.85 for all categories with all trained algorithms, except for RT181 because it could not be separated from RT027. With the models that yielded best results (RF and Light-GBM) area under the ROC curves was greater than 0.9 for RT027, RT078 and the category “Other RTs” (Figure 4). These results for AUROC curves imply that all categories but RT181 can be differentiated with high accuracy from the rest of categories.

**Figure 4.**
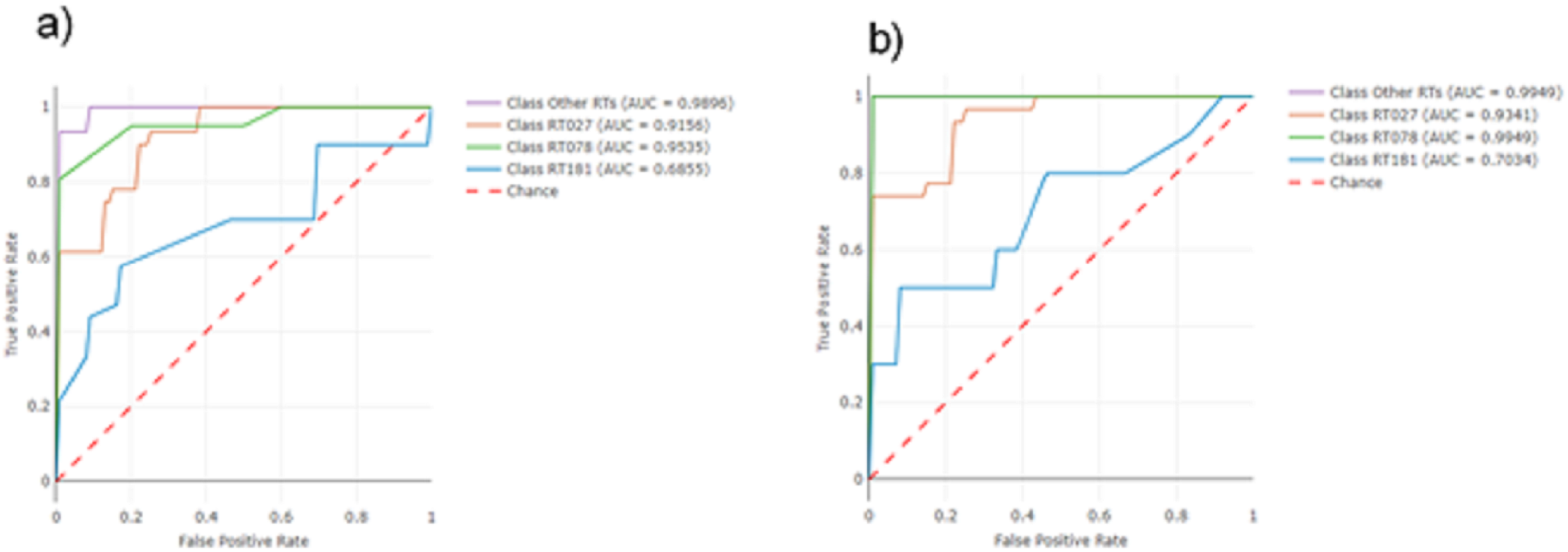
Area Under the Receiver Operating Characteristic (AUROC) curve for: a) Light-Gradient Boosting Machine algorithm; and b) Random Forest algorithm.

RT078 could be differentiated from all other RTs thanks to three putative biomarker peaks (19,222 *m/z*, 33,562 *m/z* and 41,253 *m/z*) present in these isolates and with higher intensities than in the rest (Figure 5), which AUROC = 1 and p-values in ANOVA analysis lower than 0.05 (4.85·e-25, 1.63·e-36 and 2.54·e-27, respectively). Whereas peak 19,222 *m/z* was the unique peak in this region, peaks 33,562 and 41,253 *m/z* appeared as a shift of peaks 33,840 and 40,722 *m/z*, respectively, which were present in other RTs.

**Figure 5.**
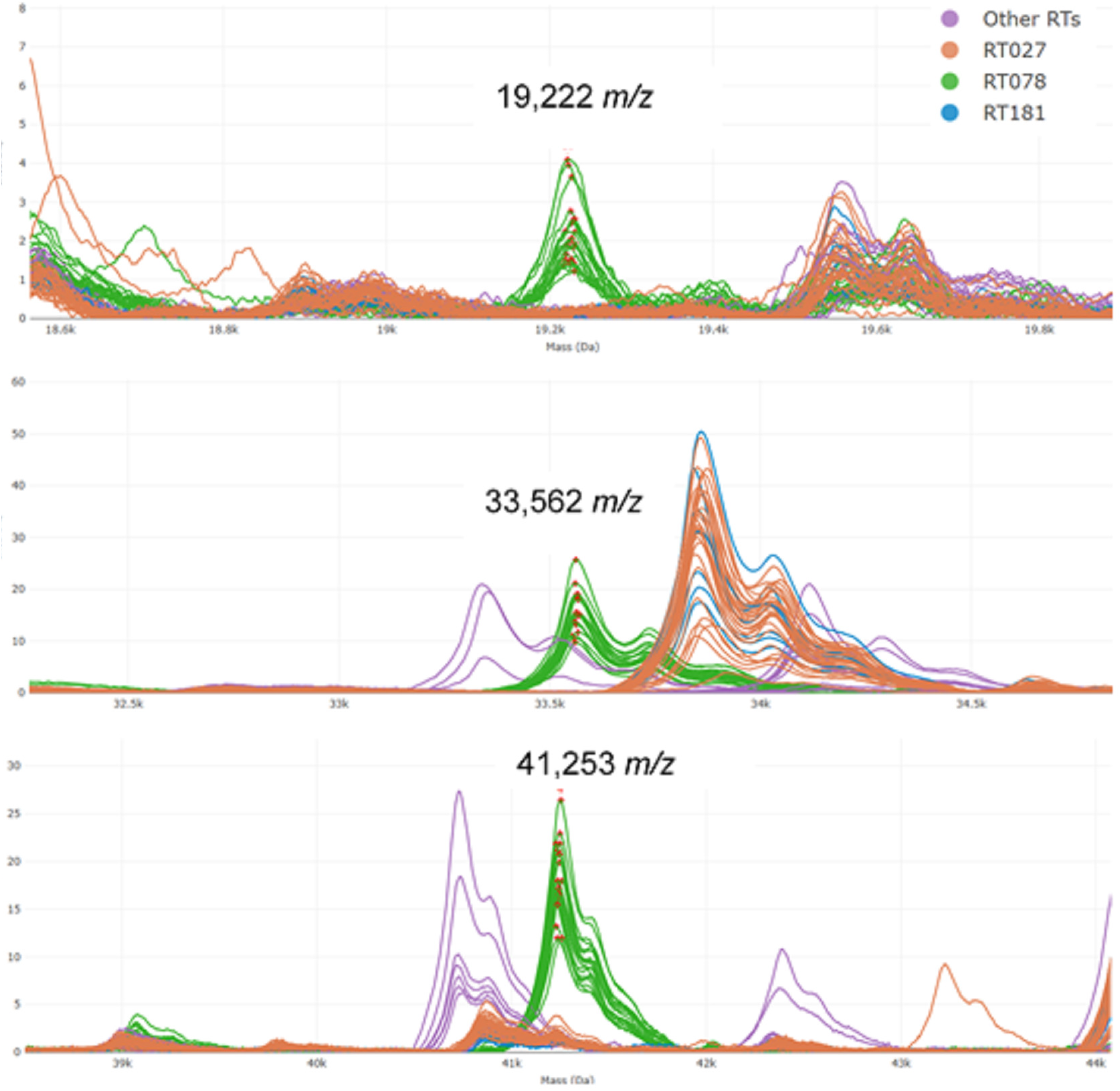
Specific biomarker peaks for the differentiation of RT078: 19,222, 22,562 and 41,253 *m/z*.

## Discussion

In this study, we evaluated the ability of MALDI-TOF MS to classify hypervirulent strains analyzing proteins with higher molecular mass than those usually evaluated for identification purposes. Our results showed that ribotyping with MALDI-TOF MS is feasible, differentiating relevant RTs like RT027 and RT078, and that this automated approach reduces the time to obtain results from several days to a few hours from the isolation of the microorganism in culture. Furthermore, the application of this technique is relatively simple, with sample preparation being identical to the preparation for routine identification using MALDI-TOF MS. Machine Learning algorithms such as RF and Light-GBM were the best to perform, both achieving a 5-fold cross validation of 88%. External validation could not be carried out due to the limited number of isolates.

The results obtained in this study with MALDI-TOF MS can compare to the molecular assay Xpert^®^ *C. difficile* BT performance, as it can separate RT027 from other clinically relevant RTs, although not from RT181. RT181 is a similar RT to RT027, as it produces toxin B, binary toxin and presents the same deletion in the regulatory gene *tcdC*. Differentiation between these two RTs can be only achievable with PCR ribotyping, which can take up to a week. The clinical implications of the differentiation between these two RTs are yet unknown as RT181 has been recently described and literature about this RT is still limited (2). Three biomarker peaks were found in our study for RT078 differentiation (19,222 *m/z*, 33,562 *m/z* and 41,253 *m/z*). Two of them, were described in a previous study (33,600±200 *m/z* and 41,375±125 *m/z*) as specific for RT078 (25). They appeared next to other peaks present in other RTs (33,840 and 40,722 *m/z*), with a shift of 300-500 *m/z*. They could correspond to different forms of the same protein, with variations between different RTs, although this was not confirmed. The peak at 19,222 *m/z* has not been described before to our knowledge. The identification of these three peaks could be applied as a fast tool for RT078 ribotyping. No other biomarkers of interest were found for the differentiation of the rest of categories.

Several studies have been published trying to ribotype *C. difficile* with MALDI-TOF MS, but the majority of them analyze the default region of identification, between 2,000 and 20,000 *m/z*, with variable results (8, 26-29). Two studies analyzed a higher molecular weight region with MALDI-TOF MS (up to 80,000 *m/z*). They achieved *C. difficile* typing by creating an internal database according to their own mass spectra (what they define the “High Molecular Weight Profile” -HMW Profile-), which then was compared to PCR ribotyping (25, 30). They could not clearly correlate their HMW profiles with conventional ribotyping, as some HMW Profiles include isolates from different conventional RTs and at the same time, some conventional RTs include isolates from different HMW profiles. They concluded that HMW ribotyping could be a useful tool for *C. difficile* outbreak detection as they detected isolates with the same HMW profile in different clinical outbreaks.

In this study *C. difficile* typing was achieved, directly correlating MALDI-TOF MS results with what is considered the Gold Standard technique in Europe, PCR ribotyping of the intergenic region between 16S rRNA and 23S rRNA. This allows for extrapolation and comparison of results with other centers that perform this technique and have a MALDI-TOF MS instrument available.

One of the flaws of this study is the limited number of isolates available, although they belong to a multicentric collection and were isolated from several hospitals in Spain. Strains were selected according to national and European epidemiology, representing the most common RTs, but were isolated only from the Spanish territory. A broader and more representative number of isolates from Europe may be needed to validate this study and confirm the results. It is possible that with an increase in the number of strains analyzed, separation of RT181 from RT027 also improves.

The methodology developed in this study is a valuable tool for the rapid discrimination of clinically relevant isolates of *C. difficile* and opens a new path for future typing studies with MALDI-TOF MS and other clinically relevant bacteria. The implementation of MALDI-TOF MS significantly reduces the time and costs for *C. difficile* ribotyping in comparison with reference methods, allowing a better optimization of hospital resources and prompt initiation of treatment according to the characterized RT, as well as more efficient and cost-effective control of the infection.

## Conflicts of Interests

MJA is employed by Clover Bioanalytical Software (Granada, Spain) but had no role in the design of the study or methodology. The rest of authors declare no conflicts of interest.

## Acknowledgments

This work is partially supported by the project PI18/00997 and PI20/00686 from the Health Research Fund (FIS. Instituto de Salud Carlos III. Plan Nacional de I+D+I 2013-2016) of the Carlos III Health Institute (ISCIII, Madrid, Spain) partially financed by the European Regional Development Fund (FEDER) ‘A way of making Europe’. The funders had no role in the study de-sign, data collection, analysis, decision to publish, or preparation/content of the manuscript. AC (Rio Hortega CM21/00165), DRT (Sara Borrell CD22/00014) and BRS (Miguel Servet CPII19/00002) are funded by ISCIII.

## Author Contributions

Conceptualization, AC, DRT, MO and BRS; Methodology, AC, MM and LA; software, MJA; formal analysis, AC and MJA; writing—original draft preparation, AC and DRT; writing—review and editing, GB, MO and BRS; supervision, MO and BRS; project administration, MO and BRS; funding acquisition, MO and BRS. All authors have read and agreed to the published version of the manuscript.

